# The Neonatal Myeloid Hypoxia Response Promotes a Cardiac Regenerative Response through Insulin-Like Growth Factor

**DOI:** 10.64898/2026.04.16.719100

**Authors:** Amanda Becker, Connor Lantz, Aparna Anathakrishman, Matthew DeBerge, Kristofor Glinton, Zhi-Dong Ge, Edward B. Thorp

## Abstract

**Background:** The adult mammalian heart lacks the regenerative potential required to replenish depleted cardiomyocytes and restore cardiac function after injury. Ischemic cardiac injury contributes to heart failure, a leading cause of death worldwide. Neonatal mice possess the capacity to regenerate injured myocardium and macrophages contribute to this process. The mechanisms contributing to the regenerative crosstalk between macrophages and cardiomyocytes remain incompletely elucidated and offer potential to inform future therapeutic strategies.

**Methods:** To test the immune contribution during cardiac regeneration, we studied the response to myocardial ischemia in neonatal mice after silencing myeloid hypoxia inducible factor 1α (*Hif1α*) and reconstituting HIF-dependent mitogens. In parallel, we examined epigenetic and transcriptional signatures of the cardiac macrophage response and focused on intercellular crosstalk with cardiomyocytes.

**Results:** In myeloid *Hif1α* deficient mice, cardiac regenerative function was lost after coronary ligation. This manifested through loss of ventricular systolic function and elevated myocardial scarring. HIF1α was found to be activated in resident-type cardiac macrophages after ischemic insult. Hypoxia stimulated macrophages to secrete insulin-like growth factor 1 (IGF-1), and this required *Hif1α*. Parallel multiomic analysis revealed epigenetic regenerative signatures.

**Conclusions:** The data reveal an age-restricted requirement for myeloid *Hif1α* in neonatal cardiac regeneration, likely through IGF-1 signaling.

## Introduction

The immune system acts to restore tissue homeostasis and healing after injury.^1^ Mechanisms of tissue healing include replacement fibrosis or tissue regeneration, and a general shift towards increased fibrotic scarring occurs with age, such as in the mammalian heart.^1^ Experimental neonatal mice, in contrast to adult mice, can regenerate injured myocardium after ischemic injury.^2^ Regenerative capacity is dependent on macrophages (Mɸs) and lost shortly after birth.^3,4,5,6^ Whether regenerative capacity is conserved in human neonates remains unclear. Case reports have described recovery of cardiac function in rare human neonates who suffer cardiac ischemic injury shortly after birth, consistent with the human capacity for cardiac regeneration cardiac, for at least a brief period after birth.^7,8^ The elucidation of underlying mechanisms of cardiac regeneration are a step towards intervening in injured hearts after the neonatal window, and towards preventing heart failure, which remains a leading cause of mortality _globally._^6,9,10,11^

During myocardial infarction (MI), impaired perfusion leads to severe tissue ischemia.^12^ Macrophages, in response to hypoxia and nutrient deprivation, stabilize hypoxia-inducible transcription factors (HIFs) to respond to the altered metabolic demands of low oxygen supply and promote cell survival.^13,14^ HIFs are a family of transcription factors with two predominate isoforms, HIF1α and HIF2α, with isoform-specific functions varying in disparate cell types.^14^ In the setting of adult murine MI, we and others have previously reported a role for myeloid HIFs in the response to cardiac injury through macrophage programming.^15,16^

Taken together, we hypothesized a role for macrophage *Hif1α* in the neonatal cardiac regeneration response. We examined regeneration after ischemic cardiac injury in neonatal mice after genetic inactivation of myeloid *Hif1α*. Through *in vivo* multiome transcriptomic analyses and complementary co-culture studies, our findings reveal macrophage HIF1α to be essential for neonatal cardiac regeneration. The underlying mechanism appears to act through transcriptional activation of insulin-like growth factor 1 (*Igf1*), and in neonatal resident-type cardiac macrophages.

## Methods

### Experimental Animals

C57BL/6J (wildtype; IMSR_JAX:000664), *LysM^cre^* (IMSR_JAX:004781), and *Hif1^flox/flox^* (IMSR_JAX:007561) mice were purchased from the Jackson Laboratory. *LysM^cre^* and *Hif1^flox/flox^* mice were backcrossed to wildtype C57BL/6J mice for 6-12 generations. To generate mice with myeloid lineage-specific deletion of *Hif1α* (*mHif1^−/−^*), *LysM^cre^* mice were bred with *Hif1^flox/flox^* mice. *Hif1^flox/flox^* mice served as littermate controls for all experimental studies unless specifically noted. Mice were housed and bred in a temperature- and humidity-controlled, pathogen free environment and kept on a 12:12 hour day/night cycle in the Northwestern Center for Comparative Medicine animal husbandry facility. Breeding partners were paired at 6-8 weeks old and had access to standard breeder chow and water *ad libitum*. Offspring from breeders were used from 0-10 days post-birth for experiments in neonates and from 8-12 weeks-old for experiments in adults. All animal studies were conducted in accordance with guidelines and regulations in an approved animal protocol by the Institutional Animal Care and Use Committee at Northwestern University (Chicago, Illinois).

### Neonatal Coronary Artery Ligation

Neonatal mice were anesthetized with 1-2% inhaled isoflurane until loss of the toe pinch reflex and then placed on an ice bed.^17,18^ Under direct microscopy, a left lateral thoracotomy was performed by making a transverse incision into the chest cavity at the anterior axillary line in the fourth intercostal space. Skin and intercostal muscles were dissected to expose the rib cage. A murine eyelid retractor was used as a rib retractor to expand the intercostal space between the third and fourth ribs and expose the anterior wall of the left ventricle. A tapered needle attached to a 10-0 non-absorbable Nylon suture (*Alcon*) was passed under the left anterior descending (LAD) coronary artery just below the branching point and tied tightly to induce infarction, as confirmed by visual blanching of the myocardium. After LAD ligation, neonates were removed from the ice bed, and the thoracic cavity was closed with a 7-0 nonabsorbable polypropylene suture (*Covidien*). The skin was then approximated and closed with tissue adhesive (SurgiSeal Stylus*, Adhezion Biomedical*). Once fully recovered from anesthesia, neonates were returned to their mothers for nursing and routine neonatal care. At reintroduction, fresh maternal urine was rubbed on neonates to help reduce cannibalization. Adult male mice were separated from pregnant females in late gestation to help reduce maternal stress and lower risk of cannibalization.

### Echocardiography

Transthoracic echocardiography was performed on mice anesthetized with isoflurane 21-days after MI surgery using the Visual Sonics Vevo 3100 equipped with a 55 MHz probe. Left ventricular systolic function was assessed in 2-dimensional M-mode with images collected 1 mm before, at, and after the papillary muscles. Measurements were made in 3 consecutive cardiac cycles and averaged for analysis. Left ventricular end-diastolic and end-systolic dimensions were measured from M-mode tracings. Fractional shortening and ejection fraction measurements were made as an indicator of systolic function using *Vevo Lab Software* (v.5.0). All echocardiography measurements were performed in a blind manner.

### Left Ventricular (LV) Infarct and Area at Risk/AAR Measurements

Neonatal mice were anesthetized with isoflurane and assessed for loss of the toe pinch reflex. Under direct microscopy, the thoracic cavity was carefully opened to maintain hemostasis and expose the heart. Using an insulin syringe with a 30-gauge needle, 100 µL FluoSpheres Polystyrene Microspheres (10 µM, red fluorescent 580/605) were injected into the left ventricle (LV). Hearts were then excised one minute later and sectioned into 1 mm coronal slices using a Mouse Heart Slicer Matrix (*Zivic Instruments*). Infarct and viable myocardium were visualized by staining each section with 1% 2,3,5-triphenyltetrazolium chloride (TTC) in saline for 15 min at 37°C and then fixing in 10% buffered formalin phosphate for 1-2 hours at 4°C. Digital infarct images were generated by placing each tissue slice directly on an Epson Perfection V600 photo scanner. Brightness and contrast were adjusted equally for each slice in ImageJ. Area at risk (AAR) was visualized by placing slices under an Olympus IX51 fluorescent scope and imaging slices with Olympus cellSens Imaging Software. Both infarct and AAR measurements were made as a percentage of the LV using ImageJ software.

### Cardiac Histology

Neonatal mice were anesthetized with isoflurane and assessed for loss of the toe pinch reflex. The thoracic cavity was carefully opened to maintain hemostasis and expose the heart which was then perfused with phosphate buffered saline (PBS) immediately followed by 4% paraformaldehyde (PFA). Hearts were then excised and placed in 10% buffered formalin phosphate for at least 12 hours overnight. After fixation, hearts were then stored in 70% ethanol, paraffin embedded, and then cross-sectioned into 10 µm thick slices. To assess gross pathology and fibrosis, sections were deparaffinized by xylene, re-hydrated, stained with Mayer’s hematoxylin for 8 minutes and then washed in running tap water for 10 minutes. Sections were then stained with picrosirius red for one hour, washed in acidified water, dehydrated, cleared in xylene, and mounted for imaging. Images were obtained at 10X to capture the entire left ventricle. ImageJ software was used to quantify fibrosis and normalized to total area of the left ventricle excluding chamber space.

### Flow Cytometry

Mice were euthanized and hearts were extensively perfused with ice-cold PBS to remove peripheral cells. Infarcted myocardium was excised, minced with fine scissors, and digested with Collagenase Type II and DNase at 37°C for 30 minutes with gentle shaking. Larger pieces of myocardial tissue were broken down with vigorous pipetting and then filtered through a 40 µm cell filter. Red blood cells (RBCs) were lysed and the number of viable cells per heart were counted on a hemocytometer with Trypan blue exclusion. Cells were then incubated with Live/Dead Zombie Aqua Fixable Dye for 15 minutes followed by an incubation period with Fc-block for 15 minutes. Cells were then labeled with fluorescently conjugated antibodies (see Ab table) for 20 minutes. To detect HIF expression in cardiac macrophages, intracellular staining was performed. Cells were fixed and permeabilized using the BD Cytofix/Cytoperm Fixation/Permeabilization Kit (*BD Biosciences*). Cells were then incubated with A647 anti-mouse HIF-1α antibody for 30 minutes. Fluorescence minus one (FMO) was used as a negative staining control. Flow cytometry was performed on a FACS LSRFortessa^TM^ X-20 cytometer (*BD Biosciences*) and data was analyzed using FlowJo software (*Tree Star*). Macrophages were identified as CD11b^+^Ly6G^−^Ly6C^LO^F4/80^+^ and were further distinguished by CD64 and Tim4. Monocytes were identified as CD11b^+^ Ly6G^−^Ly6C^HI^F4/80^LO^CD64^−^. Neutrophils were identified as CD11b^+^Ly6G^+^. Full gating strategies are depicted in figures.

### Neonatal Mouse Bone Marrow-Derived Macrophage/BMDM Isolation

Bone marrow cells were harvested from the tibia and femurs of 5-8 neonatal mice on postnatal day 1 (P1) for *in vitro* studies.^17^ Cells were released from bone using a mortar and pestle and then passed through a 70 µm filter to remove bone fragments and debris. RBCs were removed by lysis (*Biolegend*) and filtered through a 40 µm filter. Bone marrow cells were cultured in non-tissue culture 10 cm Petri dishes (*Fisherbrand)* for 7-days with Dulbecco’s Modified Eagle’s Medium (DMEM) containing 20% L929-cell conditioned media, 10% fetal bovine serum (FBS), 1% penicillin-streptomycin, 1% sodium pyruvate, and 1% L-glutamine. A half media change was performed on day 3 of culture. On day 7, adherent BMDMs were washed with PBS and 1 mL of cold Cellstripper (*Corning)* was added to the dish. After 10 minutes of incubation at room temperature, macrophages were collected, washed with DMEM medium containing 10% FBS, 1% penicillin-streptomycin, counted, and seeded to tissue culture-treated plates for experiments.

### Neonatal Mouse Cardiomyocyte Isolation

Neonatal cardiomyocytes were obtained from PND0-1 C57BL/6J mice following manufacturer protocol using the Pierce Primary Cardiomyocyte Isolation Kit (*Thermo Scientific*).^17^ Harvested hearts were digested in proprietary enzyme mixes after which single-cell suspensions of cardiac cells were plated in DMEM (*Themo Scientific*) with 10% FBS and 1% penicillin-streptomycin onto gelatin-coated 48-wll plates at a seeding density of 2.5 x 10^5^ cells/cm^2^. After 24 hours, plating media was replaced with Complete DMEM containing cardiomyocyte growth supplement. Spontaneous beating of neonatal cardiomyocytes was observed within 3 days of isolation. For proliferation experiments, neonatal cardiomyocytes were cultured in serum-starved media for 12 hours overnight followed by co-culture treatments as described below.

### Hypoxia Exposure

BMDMs were seeded in 6-well tissue culture-treated plates at a density of 3 x 10^5^ cells/well in 1 mL of DMEM containing 10% FBS and 1% penicillin-streptomycin and allowed to adhere overnight. BMDMS were then placed in a temperature- and humidity-controlled hypoxic chamber (Coy O_2_ Control In Vitro Glove Box, *Coy Laboratory Products*) and exposed to 1% O_2_ (1% O_2_, 94% N_2_, and 5% CO_2_) for 1, 3, 6, 9, 12, or 24 hours. All assays performed on BMDMs as well as BMDM supernatant were performed within the hypoxia chamber to minimize reoxygenation effects. BMDMs cultured in a temperature- and humidity-controlled tissue culture incubator supplied with room air (21% O_2_) and 5% CO_2_ were used as normoxic controls.

### Macrophage Supernatant and Cardiomyocyte Co-Culture

Neonatal cardiomyocytes were isolated and cultured in serum-starved media for 12 hours overnight followed by co-culture with hypoxic (1% O_2_ for 24 hours) or normoxic (21% O_2_ for 24 hours) neonatal BMDM supernatant for 12 hours. Immunofluorescence microscopy was then performed to evaluate cardiomyocyte proliferation.

### Immunofluorescence Microscopy of Cardiomyocytes

After co-culture, neonatal cardiomyocytes were fixed with 4% PFA for 10 minutes. Cardiomyocytes were then permeabilized with PBS + 3% Triton X100 for 10 minutes. Cells were then blocked with PBS + 0.1% Triton X100 and 3% bovine serum albumin for 1 hour, followed by incubation with primary antibodies in blocking buffer overnight. Cardiomyocytes were then washed and incubated with secondary antibodies, followed by DAPI. After washing, cardiomyocytes were imaged at 10X and 20X magnification using an Olympus Fluorescent Microscope. For each replicate, 5 distinct fields were captured. The number of positively stained nuclei were scored blindly using ImageJ (NIH) and averaged for each biological replicate.

### IGF-1 Assays

Neonatal BMDMs were subjected to hypoxic (1% O_2_) or normoxic (21% O_2_) conditions for 24 hours. Hypoxic versus normoxic BMDM supernatant was then co-cultured with neonatal cardiomyocytes for 12 hours with the addition of 0.1 µg/mL neutralizing IGF-1 inhibitor (*Abcam*) versus 0.1 µg/mL IgG followed by immunofluorescence microscopy to examine cardiomyocyte proliferation. To determine if hypoxia induces *Igf1* transcription in neonatal BMDMs, quantitative PCR was performed after 1, 3, 6, 9, 12, and 24 hours of hypoxic (1% O_2_) versus normoxic (21% O_2_) BMDM stimulation. Conditioned media was assayed at the aforementioned timepoints for IGF-1 by ELISA according to manufacturer protocol (Mouse/Rat IGF-1 DuoSet ELISA Kit, *R&D Systems*). To interrogate if HIF1α transcriptionally activates *Igf1*, ChIP PCR was performed on neonatal BMDMs after 6 hours of hypoxic (1% O_2_) versus normoxic (21% O_2_) stimulation. Flow cytometry was performed to measure IGF-1 expression in neonatal cardiac macrophage populations at days 1, 3, and 7 post-MI in wildtype neonatal mice.

### Quantitative real-time PCR

RNA was extracted from macrophages using Trizol according to the manufacturer’s instructions. Briefly, 1 mL of Trizol was added to macrophages in a 6-well plate and gently pipetted to lyse the cells. RNA was measured using a Synergy LX plate reader (*BioTek*) and 1 µg of RNA was transcribed into cDNA using the iScript cDNA Synthesis Kit (*Applied Biosciences*). Quantitative PCR was performed using the primers listed in the major resources table and SYBR Green Master Mix (*Applied Biosciences*) on a QuantStudio 3 Real-Time PCR System (*ThermoFisher*). Results are expressed in ΔΔCt values normalized to β2m.

### Chromatin Immunoprecipitation/ChIP and qPCR

*In-silico* interrogation of the mouse genome was performed using the UCSC Genome Browser to identify potential hypoxia response element (HRE) sequences in promoter and enhancer regions on *Igf1* (chromosome 10). Please refer to the major resources table for generated forward and reverse primers based on our *in-silico* analysis. The Abcam High-Sensitivity ChIP Kit was utilized according to the manufacturer’s protocol to isolate macrophage DNA, perform HIF1α antibody (*Novus Biologicals*) immunoprecipitation, clean HIF1α/DNA complexes, reverse cross linking, and purify DNA for PCR. Real time PCR analysis was performed using the primers listed in the key resources table, SYBR Green Master Mix (*Applied Biosciences*), and 2 µL of eluted DNA on a QuantStudio 3 Real-Time PCR System (*ThermoFisher*). Fold enrichment was calculated normalized to the Abcam kit’s non-immune IgG.

### Multiome Single Nuclei RNA Sequencing and Single Nuclei ATAC Sequencing Nuclei Isolation

Neonatal coronary artery ligation was performed on P1 (regenerative-window) and P10 (non-regenerative window) wildtype C57BL/6J neonatal mice. Seven days after MI, mice were euthanized and hearts were extensively perfused with ice-cold PBS to remove peripheral cells. Infarcted left ventricles were then excised and flash frozen and stored in liquid nitrogen. The 10X Genomics Chromium Nuclei Isolation Kit (PN-1000494) was used to isolate cardiac nuclei in direct accordance with the manufacturer’s protocol. Isolated nuclei were then delivered to the Northwestern University Sequencing Core for library preparation and sequencing.

### Multiome single nuclei snRNA-Sequencing and snATAC-Sequencing

Nuclei number was analyzed using the Nexcelom Cellometer Auto2000 with AOPI fluorescent staining method. Nuclei then underwent transposition with ATAC enzyme for one hour at 37°C. Sixteen-thousand transposed nuclei were then loaded into the Chromium Controller (*10X Genomics*, PN-120223) on a Chromium Next GEM Chip J (*10X Genomics*, PN-1000230) and processed to generate single cell gel beads in the emulsion (GEM) according to the manufacturer’s protocol. Barcoded DNA and cDNA were PCR amplified and subjected to library construction. The single nuclei ATAC-seq library were generated using the Chromium Next GEM Single Cell Multiome ATAC + Gene expression kit (*10X Genomics*, PN-1000281) and single Index Kit N Set A (*10X Genomics*, PN-1000212) according to the manufacturer’s manual. In addition, the amplified cDNA was used for gene expression library using dual Index Kit TT Set A (*10X Genomics*, PN-1000215). Quality control for the constructed library was performed by Agilent Bioanalyzer High Sensitivity DNA kit (*Agilent Technologies*, 5067-4626) and Qubit DNA HS assay kit for qualitative and quantitative analysis, respectively. For the snATAC-seq library, the multiplexed libraries were pooled and sequenced on Illumina Novaseq X Plus sequencer with 100 cycles kits using the following read length: 50 bp Read1 and 49 bp Read2. For snRNA-seq library, the libraries were sequenced on Illumina Novaseq sequencer with 100 cycles kits using the following read length: 28 bp Read 1 for cell barcode and UMI and 90 bp Read 2 for transcript expression. The targeted sequencing depth for snATAC-seq and snRNA-seq was 25,000 and 20,000 reads per cell, respectively.

### Single-nucleus Multiome Sequencing and Data Processing

Single-nucleus RNA and ATAC sequencing was performed using the 10x Genomics Multiome platform on nuclei isolated from cardiac tissue of P1 and P10 mice following myocardial infarction. Raw sequencing data were aligned to the GRCm39 (mm39) reference genome using Cell Ranger ARC (v2.1.0) with BAM file output enabled and otherwise default parameters. Peak calling for the ATAC assay was performed using MACS2, and a unified peak set was generated across all samples to enable cross-sample comparisons. Data were analyzed in R using Seurat (v5.4.0) and Signac. Nuclei were filtered to retain those with 500–7,500 RNA features, mitochondrial read fraction below 30%, 500–30,000 ATAC features, TSS enrichment score greater than 1, and nucleosome signal below 2. RNA counts were normalized using NormalizeData and scaled using ScaleData. For the ATAC assay, term frequency-inverse document frequency (TF-IDF) normalization was applied, followed by singular value decomposition (SVD) via RunSVD for linear dimensionality reduction. Batch effects across samples were corrected for both RNA and ATAC modalities using Harmony (v1.2.4). Weighted nearest neighbor (WNN) analysis was performed integrating the Harmony-corrected RNA and ATAC embeddings to generate a joint UMAP representation. Unsupervised clustering was performed on the WNN graph, and clusters were annotated based on the expression of canonical marker genes and cell type enrichment analysis using Enrichr. Myeloid cells were isolated and re-clustered independently. Subpopulations were defined by the expression of lineage marker genes and by module scoring using gene sets associated with cardiac-resident and recruited macrophage populations.

### Chromatin Accessibility and Transcription Factor Activity Analysis

ChromVAR (v.1.28.0) was used to compute transcription factor motif accessibility scores using the JASPAR2020 motif database. Activity of the ARNT::HIF1A transcription factor complex was specifically examined within macrophage populations. Chromatin accessibility at genomic loci of interest was visualized using CoveragePlot, and gene expression was visualized using violin plots. To capture a broader set of candidates for downstream validation, differentially expressed genes between conditions were identified using the Wilcoxon rank-sum test with a p-value threshold of 0.1 and a minimum average log2 fold change of 0.2. P-values were adjusted for multiple comparisons using the Benjamini–Hochberg method as well to determine statistically significant genes.

### Processing and Analysis of Single Cell mRNA Sequencing Data

All data analyses were performed in direct accordance with our previously published methodologies.^17^ Please refer to our previous publication for detailed methodology regarding the following: Raw Sequencing Data Processing (Alignment, Barcode Assignment, and UMI Counting), Filtering and Normalization, Unsupervised Clustering, Cluster Annotation, Sub-setting Myeloid Cells, Differential Gene Expression, Gene Ontology Enrichment, and Cell Communication.^17^ A detailed description of our code and analysis is available on GitHub (https://github.com/thorplab).

### Quantification and Statistical Analyses

Statistical analyses were performed with GraphPad Prism 9 software. Comparisons between two groups were performed using a two-tailed, Mann-Whitney *t* test with a 95% confidence interval. For comparisons of more than two variables, a one-way or two-way ANOVA was used with a 95% confidence interval; when necessary, a Tukey test was used to correct for multiple comparisons. For *in vivo* experiments, sample size is indicated in figures/legends and represent pooled data from two or more independent experiments. For experiments in culture, experimental sample size is indicated in figures/legends and are representative of data from two or more independent experiments. All data are presented as ± SEM. Criteria for significant differences (*, P < 0.05; **, P < 0.01; ***, P < 0.001; ****, P < 0.0001) are located in the figure legends. Analyses labeled “ns” are not statistically significant.

#### Major Resources Table

Please see table in Supplemental Materials.

#### Resource Availability and Lead Contact

Further information and requests for resources and reagents should be directed to Amanda Becker or Edward B. Thorp.

#### Materials Availability

All reagents generated in this study are available upon reasonable request from the lead contact without restriction.

#### Data and Code Availability

The multiome scRNA-seq and scATAC-seq datasets used in this study are deposited in GEO and are already publicly available. The accession number is listed in the Major Resources Table. Microscopy data reported in this paper will be shared by the lead contact upon request. All original code was deposited on GitHub (https://github.com/thorplab) and is already publicly available. DOIs are listed in the Major Resources Table. Any additional information required to reanalyze the data reported in this paper is available from the lead contact upon request.

#### Independent Data Access and Analysis

Dr. Becker and Thorp had full access to all of the data in this study and takes responsibility for its integrity and analysis. Dr. Lantz and Thorp had full access to all of the neonatal multiome data and scRNA-sequencing data in the study and takes responsibility for its integrity and analysis.

## Results

### Neonatal cardiac *C1q*^+^TLF^+^ (*Timd4, Lyve1, and Folr2*) macrophages remodel chromatin and mobilize HIF1α-dependent pathways in response to ischemic injury

To ascertain key regulatory nodes in the neonatal immune response to injury, we initially examined the cellular innate immune response in C57BL/6J neonatal mice at timepoints following permanent coronary artery ligation on postnatal day 1 (P1). Cardiac neutrophils (defined as CD11b^+^Ly6G^+^)^15,17^ were monitored (**Figures S1A, S1B**), as well as monocytes (CD11b^+^Ly6G^−^Ly6C^+^). Monocytes preferentially accumulated within the injured myocardium on day 1 with subsequent return to steady state by day 7 post-MI (**Figure S1B**). In contrast to neutrophils and monocytes, the number of macrophages (CD11b^+^Ly6G^−^Ly6C^−^CD64^+^) steadily increased up to 7 days after MI (**Figure S1B**). Therefore, we selected this temporal window to assess differences in macrophage transcriptional signatures and chromatin accessibility in regenerative versus non-regenerative neonatal hearts.

At 7 days post-MI, we performed multiome single nuclei RNA-sequencing (snRNA-seq) and single nuclei ATAC-sequencing (snATAC-seq) on infarcted left ventricles from regenerative (P1) versus non-regenerative (P10) C57BL/6J wildtype neonatal mice. Previous studies have identified that murine cardiac regenerative capacity is lost within the first week of life.^4,5,6^ Thus, P10 neonates no longer possess regenerative potential. Weighted Nearest Neighbor (WNN) analysis was performed on snRNA-seq combined with snATAC-seq to identify cardiac cells populations 7-days post-MI (**Figures 1A, S2A**). Both P1 and P10 neonatal hearts were found to have increased resident cardiac macrophages compared to recruited macrophages (**Figure 1B**). Next, we sub-clustered resident and recruited macrophages by examining differences in chromatin accessibility at key markers of resident (*Mertk*) and recruited (*Ccr2*) macrophages as well as scoring previously described gene modules and macrophage canonical marker expression to better resolve differences in transcriptional heterogeneity (**Figures 1C, S2B**).^17^ We then performed gene ontology enrichment analysis to glean insight into differences in P1 versus P10 resident cardiac macrophage function. P1 resident cardiac macrophages upregulated pathways associated with organ development, cell cycle, cell migration, chromatin remodeling, and muscle tissue development (**Figure 1D**). In comparison, P10 resident cardiac macrophages selectively upregulated pathways involved in oxidative phosphorylation, response to oxygen levels and hypoxia, retrograde endocannabinoid signaling, cytoprotection by HMOX1, and oxygen transport (**Figure 1E**).

**Figure 1:**
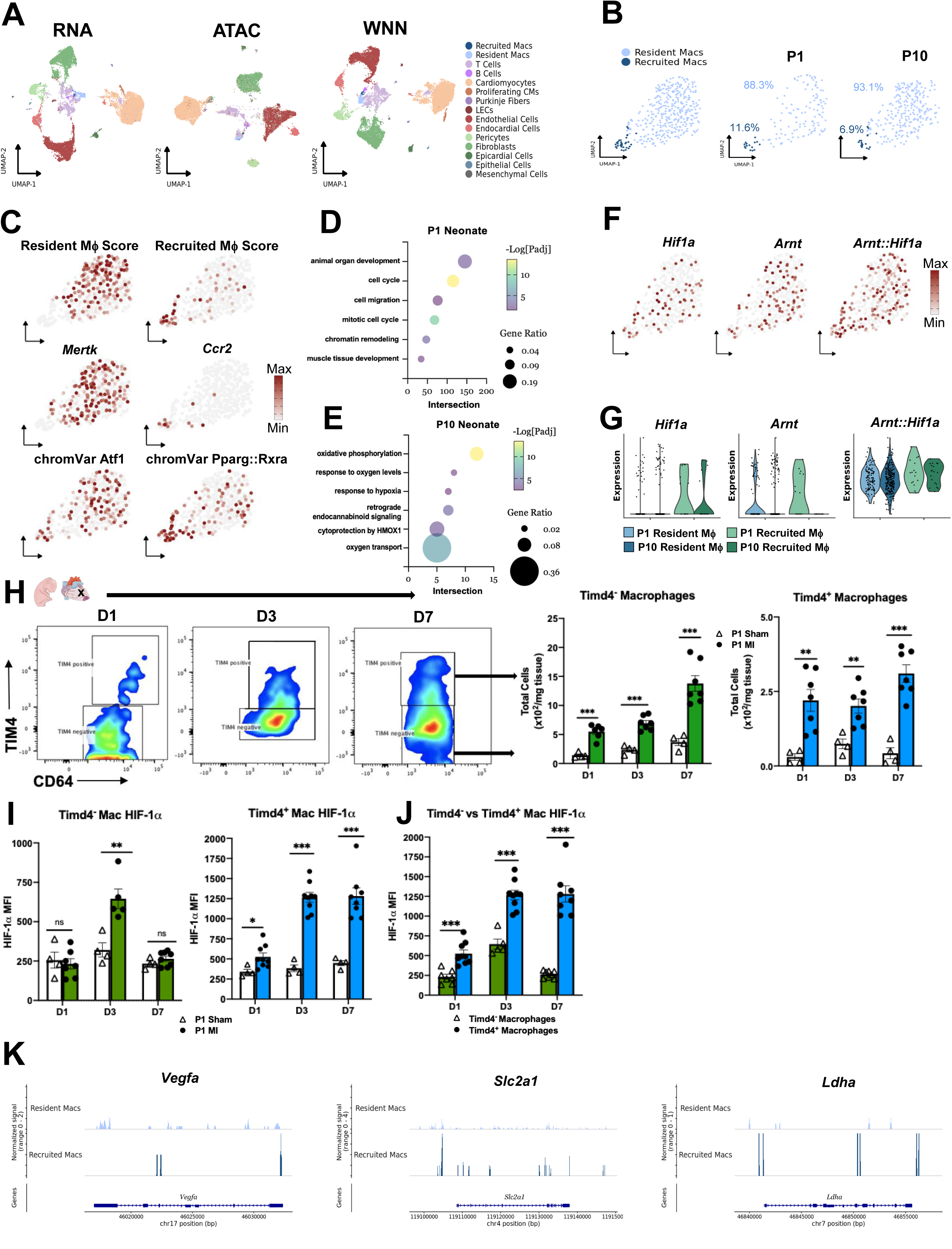
snRNA-seq combined with snATAC-seq revealed distinct neonatal cardiac macrophage populations that respond differently to decreased oxygen levels after ischemic cardiac injury. A. Weighted Nearest Neighbor analysis was computed on snRNA-seq combined with snATAC-seq to identify cardiac cell types in P1 and P10 C57BL/6 wildtype neonatal mice 7 days post-MI (total cells= 16,582 cells; *n*= 3 mice/group). B. Representative proportions of resident and recruited macrophages. C. Sub-clustering of resident and recruited macrophages was determined by scoring previously described gene modules and macrophage canonical marker expression.^17^ D. Top differentially expressed gene features in P1 resident cardiac macrophages 7 days post-MI. E. Top differentially expressed gene features in P10 resident cardiac macrophages 7 days post-MI. F. Expression of *Hif1α*, *Arnt* (*Hif1β*) and *ARNT::Hif1α* motifs determined by chromVAR in resident and recruited neonatal macrophages 7 days post-MI. G. Expression of *Hif1α*, *Arnt* (*Hif1β*) and *ARNT::Hif1α* motifs across P1 and P10 resident and recruited macrophages. H. Quantification of TIM4^+^ and TIM4^−^ macrophages by flow cytometry in wildtype neonatal mice at Days 1, 3, and 7 post-MI (*n*= 4-7). I. HIF1α protein expression from TIM4^+^ and TIM4^−^ macrophages in wildtype neonatal mice at Days 1, 3, and 7 post-MI (*n*= 4-8). J. Difference in HIF1α protein expression from TIM4^+^ vs TIM4^−^ macrophages in wildtype neonatal mice at Days 1, 3, and 7 post-MI (*n*= 5-9). K. Differences in chromatin organization around canonical HIF1α target genes, *Vegfa, Slc2a1, and Ldha* in recruited vs resident cardiac macrophages 7 days post-MI. Each data point represents a biological replicate from an individual mouse, and data are representative of experiments repeated at least three times. Data with error bars are presented as mean ± SEM. NS= not significant, *, *p* <0.05, ** *p* <0.01, *** *p* <0.001, **** *p* <0.0001.

Next, we examined differences in *Hif1α* expression between P1 and P10 resident versus recruited macrophages. Both P1 and P10 resident cardiac macrophages had increased *Hif1α* expression compared to recruited macrophages (**Figure 1G**). To validate neonatal resident cardiac macrophage HIF1α levels in *vivo*, we first examined resident (Timd4^+^) versus non-resident macrophage (Timd4^−^) accumulation within infarcted left ventricles from C57BL/6J wildtype neonatal mice on days 1, 3, and 7 following MI on P1. Consistent with general macrophage accumulation post-MI (**Figure S1B**), non-resident macrophages steadily increased in the left ventricle up to 7 days after MI (**Figure 1H**). Resident macrophages expanded on day 1 and remained increased in number through day 7 post-MI (**Figure 1H**). Next, we measured HIF1α accumulation in Timd4^−^and Timd4^+^ macrophages on days 1, 3, and 7 post-MI (**Figures 1I, S1C, S1D**). We observed increased HIF1α accumulation in Timd4^+^ compared to Timd4^−^ macrophages at all time points, with HIF1α accumulation beginning on day 1, peaking at day 3, and remaining consistently increased at day 7 post-MI (**Figure 1J**).

As an independent corroboration of *Hif1α* expression in P1 resident cardiac macrophages, we turned to our previously published single-cell mRNA sequencing (scRNA-seq) dataset, in which scRNA-seq was performed 7 days post-MI on enriched CD45^+^ cells from neonatal (P1) and adult C57BL/6J wildtype mice, to interrogate where *Hif1α* expression was present in myeloid cells. Uniform manifold approximation and projection representation of our previously published myeloid cell clusters (**Figure S2C**) and the percentage of transcriptionally distinct macrophage populations in neonatal versus adult mice (**Figure S2D**) post-MI, necessary for subsequent *Hif1α* interrogation, are provided for reference.^17^ In P1 neonatal mice, *Hif1α* was expressed at steady state and enriched post-MI in C1q^+^TLF^+^ resident cardiac macrophages (**Figure S2E**). Interestingly, loss of *Hif1α* enrichment was seen in adult *C1q*^+^TLF^+^ resident cardiac macrophages post-MI (**Figure S2E**). Gene ontology analysis of P1 neonatal *C1q*^+^TLF^+^ resident cardiac macrophages revealed upregulated pathways associated with cardiac development, angiogenesis, regeneration, cellular responses to hypoxia, and insulin signaling (**Figures S2H**). Contrastingly, adult C1q^+^TLF^+^ macrophages upregulated pathways involved in antigen presentation, inflammation, myeloid leukocyte migration, programmed cell death, and hydroxylation of HIF-1α (**Figure S2I**).

We next examined differences in chromatin organization around known canonical HIF1α targets from the ATAC dataset. We measured distinct signatures in chromatin accessibility for *Vegfa*, *Slc2a1*, and *Ldha* between neonatal resident and recruited macrophages after MI (**Figure 1K**). These data implicate disparate HIF1α and age-specific macrophage functions in response to ischemic injury, with changes in macrophage function beginning in the first week of life.

### Myeloid *Hif1α* is necessary for neonatal cardiac regeneration after ischemic cardiac injury

To assess the causal contribution of myeloid *Hif1α* in cardiac regeneration, we subjected myeloid *Hif1α* expressing mice (referred to as *mHif1^+/+^*) and mice with genetic depletion of myeloid *Hif1α* via the *LysMCre* system (referred to as *mHif1^−/−^*) to permanent ligation of the left anterior descending (LAD) coronary artery on P1 and measured indices of cardiac injury, including left ventricular function and histologic morphology, to assess impact on regeneration (**Figure 2A**). Loss of *Hif1α* exacerbated initial cardiac injury as seen by larger left ventricular infarct sizes and areas at risk (AAR) 7 days post-MI (**Figure 2B**) compared to *mHif1^+/+^* neonatal mice. Assessment of fibrotic scaring by picrosirius red staining 21 days post-MI revealed significant myocardial collagen deposition in *mHif1^−/−^*mice compared to *mHif1^+/+^* mice whose left ventricles showed minimal evidence of scarring (**Figure 2C**). Additionally, echocardiography performed 21 days post-MI revealed significantly impaired left ventricular function as evidenced by decreased ejection fraction and fractional shortening as well as reduced left ventricular anterior wall thickness during systole and increased left ventricular volume during both systole and diastole in *mHif1^−/−^*neonatal mice compared to *Hif1α* floxed littermate controls (**Figure 2D**). Interestingly, heterozygous LysMCre (*mHif1^+/−^*) neonatal mice also had significantly increased left ventricular scarring (**Figure 2C**) and impaired left ventricular function (**Figure 2D**) 21 days post-MI compared to *Hif1α* floxed littermates suggesting that a gene dosage reduction in *Hif1α* contributes to disparate phenotypes. Together, our findings implicate the necessity of *Hif1α* in neonatal cardiac regeneration after ischemic injury.

**Figure 2:**
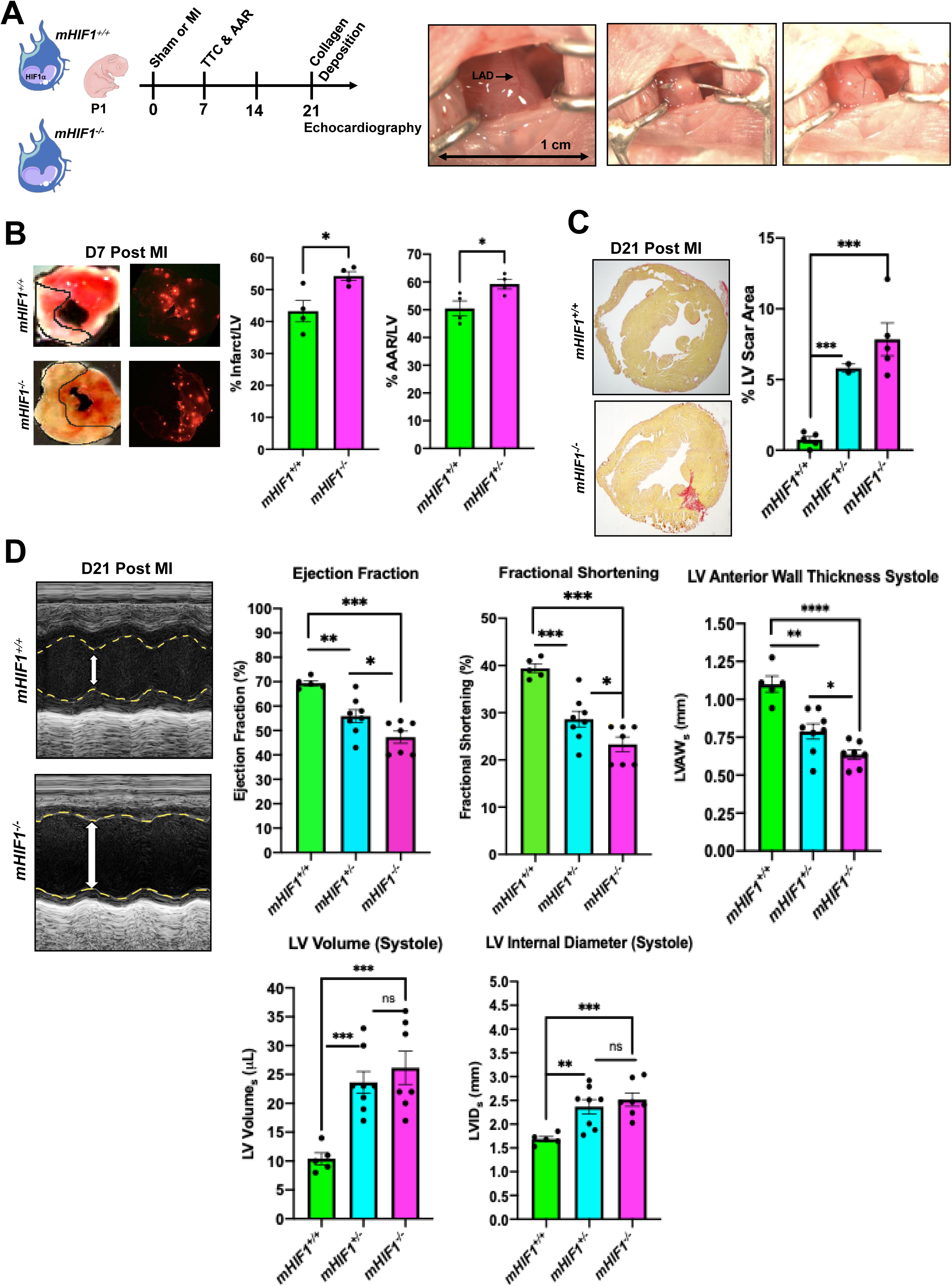
Myeloid *Hif1α* is necessary for neonatal cardiac regeneration after ischemic cardiac injury. A. Control (referred to as *mHif1^+/+^*) and genetic macrophage depletion of *Hif1α* (referred to as *mHif1^−/−^*) neonatal mice were subjected to permanent LAD coronary artery ligation to induce MI on postnatal day 1 (P1). Cardiac function assessed by echocardiography and histology were performed at specifics time points post-MI. B. Representative 1 mm cardiac sections from *mHif1^+/+^* and *mHif1^−/−^* 7 days post-MI stained with triphenyltetrazolium chloride (TTC) for infarct area measurements or injected with fluorescent microspheres to quantify the area at risk (AAR). The ratio of % area infarct size of the LV and % AAR of the LV were quantified (*n* = 4). C. Assessment of fibrotic scaring using picrosirius red staining of collagen deposition at the site of myocardial injury 21 days post-MI in *mHif1^+/+^* and *mHif1^−/−^*neonatal mice (*n* = 5). D. Quantification of ejection fraction (EF), fractional shortening (FS), left ventricle anterior wall thickness during systole (LVAW_s_), left ventricle volume during systole, and left ventricle internal diameter during systole (LVID_s_) 21 days post-MI in *mHif1^+/+^* and *mHif1^−/−^* neonatal mice (*n* = 5-8). Each data point represents a biological replicate from an individual mouse, and data are representative of experiments repeated at least two times. Data with error bars are presented as mean ± SEM. NS= not significant, *, *p* <0.05, ** *p* <0.01, *** *p* <0.001, **** *p* <0.0001.

### Hypoxia stimulates neonatal HIF1α^+^ macrophages to secrete IGF-1

To test the hypothesis that hypoxia stimulates neonatal macrophages to secrete factors that induce cardiomyocyte proliferation, we harvested BMDMs from P1 C57BL/6J neonatal mice followed by hypoxic (1% O_2_) versus normoxic (21% O_2_) conditioning for 24 hours. We then co-cultured hypoxic or normoxic macrophage supernatant with neonatal cardiomyocytes harvested from P1 C57BL/6J mice for 12 hours followed by immunofluorescent staining to assess cardiomyocyte proliferation (**Figure 3A**). Supernatant from hypoxic neonatal macrophages induced markers of cardiomyocyte proliferation compared to co-culture of cardiomyocytes with supernatant from normoxic neonatal macrophages (**Figure 3B**). To assess the necessity of HIF1α in macrophage mitogen production, we harvested BMDMs from P1 *mHif1^−/−^* neonatal mice followed by hypoxic versus normoxic conditioning and then co-culture as previously stated. With *Hif1α* deficiency, hypoxic neonatal macrophages no longer secreted mitogens that induced cardiomyocyte proliferation (**Figure 3C**). Of note, *Hif1α*^+^ and *Hif1α*^−^ macrophages had similar survival with hypoxic stimulation (**Figure 3A**).

**Figure 3:**
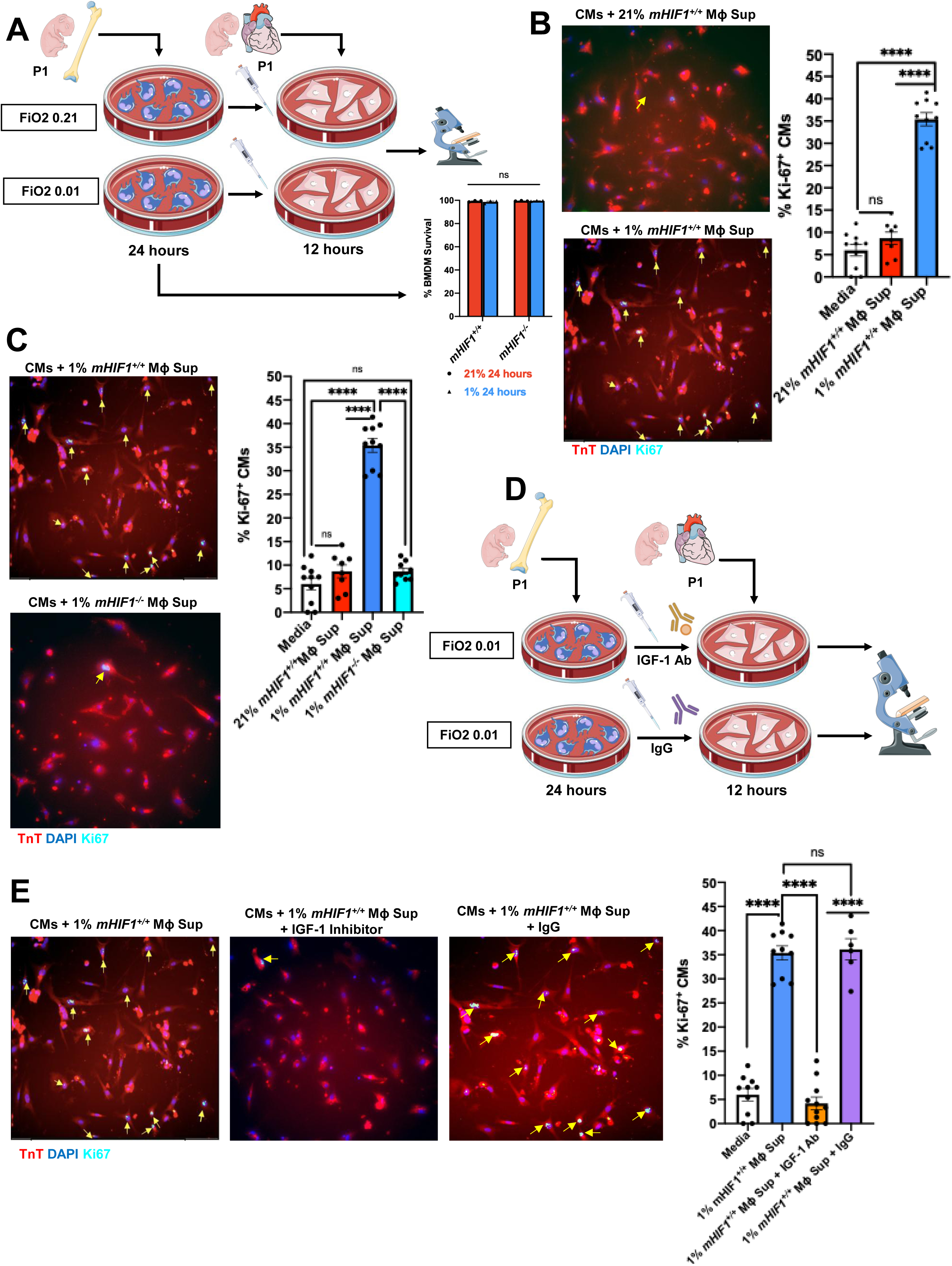
Hypoxia stimulates neonatal *Hif1α^+^* macrophages to secrete IGF-1, a cardiomyocyte mitogen. A. Neonatal bone marrow-derived macrophages (BMDMs) harvested from wildtype mice on P1 were subjected to hypoxic (FiO_2_ 0.01) or normoxic (FiO_2_ 0.21) conditions for 24 hours followed by assessment of BMDM survival via flow cytometry with Zombie Live/Dead staining (*n* = 6-8 mice). BMDM supernatant was then co-cultured with wildtype neonatal P1 cardiomyocytes for 12 hours followed by assessment of cardiomyocyte proliferation. B. Proliferation of neonatal cardiomyocytes (defined by positive nuclear Ki67^+^ expression) cultured with supernatant from hypoxic vs normoxic *mHif1^+/+^* BMDMs (*n* = 6-8 mice; scale bar 100 um). C. Proliferation of neonatal cardiomyocytes cultured with supernatant from hypoxic *mHif1^+/+^* vs *mHif1^−/−^* BMDMs (*n* = 6-8 mice; scale bar 100 um). D. Neonatal BMDMs harvested from wildtype mice on P1 were subjected to hypoxic (FiO_2_ 0.01) conditions for 24 hours. BMDM supernatant was then co-cultured with neonatal P1 cardiomyocytes for 12 hours with the addition of 0.1 ug/mL neutralizing IGF-1 inhibitor vs 0.1 ug/mL IgG followed by assessment of cardiomyocyte proliferation. E. Proliferation of neonatal cardiomyocytes cultured with supernatant from hypoxic *mHif1^+/+^* BMDMs and neutralizing IGF-1 inhibitor vs non-specific IgG (*n* = 6-8 mice; scale bar 100 um). Each data point represents a set of cells, and data are representative of experiments repeated at least two times. Data with error bars are presented as mean ± SEM. NS= not significant, *, *p* <0.05, ** *p* <0.01, *** *p* <0.001, **** *p* <0.0001.

Based on our evaluation of the top differentially expressed genes from neonatal *C1q*^+^TLF^+^ resident cardiac macrophages post-MI (**Figures S2F, S2G**), we hypothesized that insulin-like growth factor 1 (IGF-1) was a potential mitogen secreted from hypoxic *Hif1α^+^* macrophages. To test this hypothesis, we added a neutralizing IGF-1 inhibitor versus non-specific IgG in co-culture of neonatal cardiomyocytes with supernatant from hypoxic neonatal HIF1α^+^ macrophages (**Figure 3D**). We measured loss of cardiomyocyte proliferation markers with the addition of a neutralizing IGF-1 inhibitor (**Figure 3E**). Cardiomyocyte proliferation was preserved with the addition of control IgG (**Figure 3E**). These data are consistent with *Hif1α* dependent secretion of IGF-1 from hypoxic P1 neonatal macrophages.

### Hypoxia stimulates *Hif1α*-dependent *Igf1* induction in neonatal macrophages

To determine the kinetics of *Igf1* induction in hypoxic neonatal macrophages, we harvested BMDMs from P1 C57BL/6J neonatal mice and measured changes in *Igf1* expression at specific timepoints with hypoxic (1% O_2_) versus normoxic (21% O_2_) conditioning. *Igf1* expression began to increase with 1 hour of hypoxic stimulation, peaked at 6 hours, and remained increased at 24 hours compared to normoxic controls (**Figure 4A**). To assess whether *Hif1α* was required for hypoxic *Igf1* induction, we harvested BMDMs from *mHif1^−/−^* neonatal mice and once again measured changes in *Igf1* expression at specific timepoints with hypoxic versus normoxic conditioning. In the absence of *Hif1α*, hypoxic *Igf1* induction was lost at all timepoints (**Figure 4A**). We then measured IGF-1 secretion from hypoxic versus normoxic neonatal *Hif1α^+^* macrophages to confirm that *Igf1* induction was accompanied by the translation and secretion of IGF-1 from hypoxic macrophages. IGF-1 release increased after 1 hour of hypoxic stimulation, peaked at 9 hours, and remained increased at 24 hours compared to normoxic *Hif1α^+^* neonatal macrophages (**Figure 4B**). To determine whether HIF1α transcriptionally activates *Igf1* in neonatal macrophages, we performed *in silico* analysis of the mouse genome to identify potential hypoxia response element (HRE) sequences for HIF1α binding on *Igf1* (**Figure S3A**). We then subjected neonatal macrophages to 6 hours of hypoxic or normoxic conditioning followed by the application of ChIP qPCR. Subsequently, we newly identify that hypoxia stimulates HIF1α−dependent mRNA accumulation of *Igf1* (**Figures 4C, S3B**). To test for *Igf1* expression *in vivo*, we measured IGF-1 protein accumulation in Timd4^−^ and Timd4^+^ neonatal *Hif1α^+^* cardiac macrophages on days 1, 3, and 7 post-MI on P1. Timd4^−^ cardiac macrophages had increased IGF-1 compared to controls on day 1 post-MI, although significantly less than Timd4^+^ macrophages, with return to baseline IGF-1 levels by day 3 post-MI (**Figures 4D**, **S3C**). Contrastingly, Timd4^+^ cardiac macrophages had robust IGF-1 accumulation at all timepoints post-MI (**Figures 4E, S3D**) and significantly more IGF-1 than Timd4^−^macrophages (**Figure 4F**). Taken together, hypoxia stimulates HIF1α mRNA accumulation of *Igf1* and subsequent P1 neonatal macrophage release of IGF-1.

**Figure 4:**
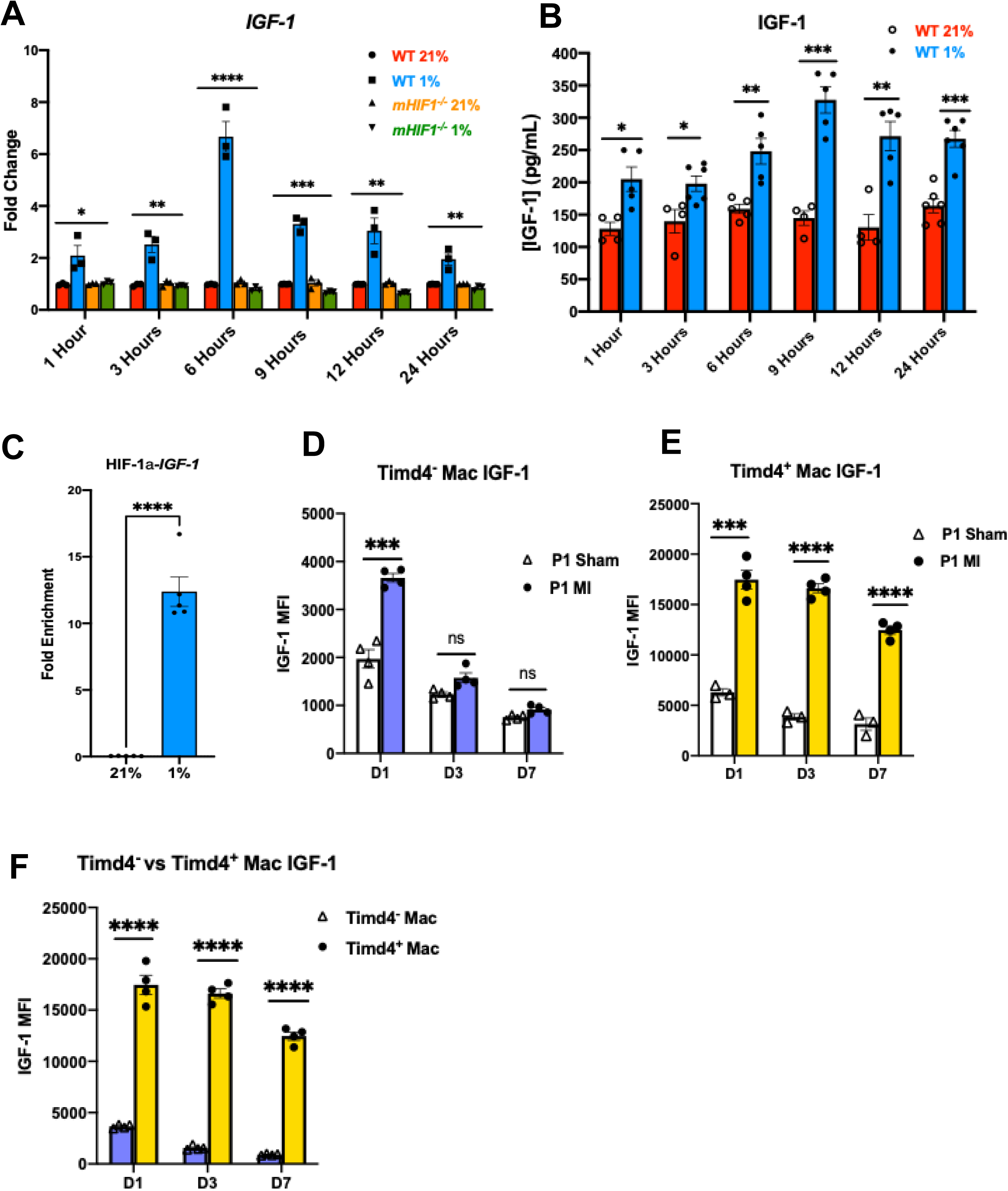
Hypoxia stimulates *Hif1α*-dependent *Igf1* induction in neonatal macrophages. A. *Igf1* induction in *Hif1α^+^* vs *Hif1α^−^* macrophages at specific timepoints with hypoxic (FiO_2_ 0.01) vs normoxic (FiO_2_ 0.21) stimulation. B. B. IGF-1 secretion from hypoxic vs normoxic *Hif1α^+^* macrophages at specific timepoints with hypoxic (FiO_2_ 0.01) vs normoxic (FiO_2_ 0.21) stimulation. C. C. Hypoxia (FiO_2_ 0.01) stimulates HIF1α binding to hypoxia response element (HRE) sites on *Igf1*. D. D. IGF-1 protein expression from TIM4^−^ cardiac macrophages in *Hif1α*^+^ wildtype neonatal mice at Days 1, 3, and 7 post-MI (*n*= 3-4). E. E. IGF-1 protein expression from TIM4^+^ cardiac macrophages in *Hif1α*^+^ wildtype neonatal mice at Days 1, 3, and 7 post-MI (*n*= 4). F. IGF-1 protein expression from TIM4^−^ vs TIM4^+^ cardiac macrophages in *Hif1α*^+^ wildtype neonatal mice at Days 1, 3, and 7 post-MI (*n*= 4). Each data point represents a biological replicate from an individual mouse, and data are representative of experiments repeated at least three times. Data with error bars are presented as mean ± SEM. NS= not significant, *, *p* <0.05, ** *p* <0.01, *** *p* <0.001, **** *p* <0.0001.

### IGF-1 signaling occurs between neonatal *C1q*^+^TLF^+^ *HIF-1α*^+^ cardiac macrophages and cardiomyocytes after MI

To gain insight into neonatal IGF-1 signaling after MI, we integrated our previously published scRNA-seq dataset with Cui et al.’s single-nuclei RNA-sequencing dataset (**Figure 5A**), which includes cardiomyocyte transcriptomic data, to perform *CellChat* analyses.^17,19^ This model allowed us to statistically infer IGF-1 ligand-pair signaling between neonatal cardiomyocytes, immune cells, vascular and endothelial cells, and stromal cells after MI (**Figures 5B, S4A**). We observed enrichment of *Igf1* in neonatal cardiac macrophages, with greatest enrichment occurring in *C1q*^+^TLF^+^ macrophages, and enrichment of *Igf1r* in neonatal cardiomyocytes (**Figure 5C**) after MI. As *Igf1* message was also enriched in *Spp1^+^* and *Arg1^+^* macrophages, we performed subsequent *CellChat* analyses to identify dominant macrophage IGF-1 senders and cardiomyocyte IGF-1 receivers during regeneration. We identified that *C1q*^+^TLF^+^ macrophages were the strongest suggested senders of IGF-1 and that type 5 cardiomyocytes were the greatest receivers of IGF-1 (**Figure 5D**). Next, we aligned *Hif1α* and *Igf1* expression in neonatal and adult *C1q*^+^TLF^+^ macrophages to examine if changes in expression occur between regenerative and non-regenerative hearts. We observed reductions in *Hif1α* and *Igf1* expression in adult compared to neonatal *C1q*^+^TLF^+^ cardiac macrophages (**Figure S4B**).

**Figure 5:**
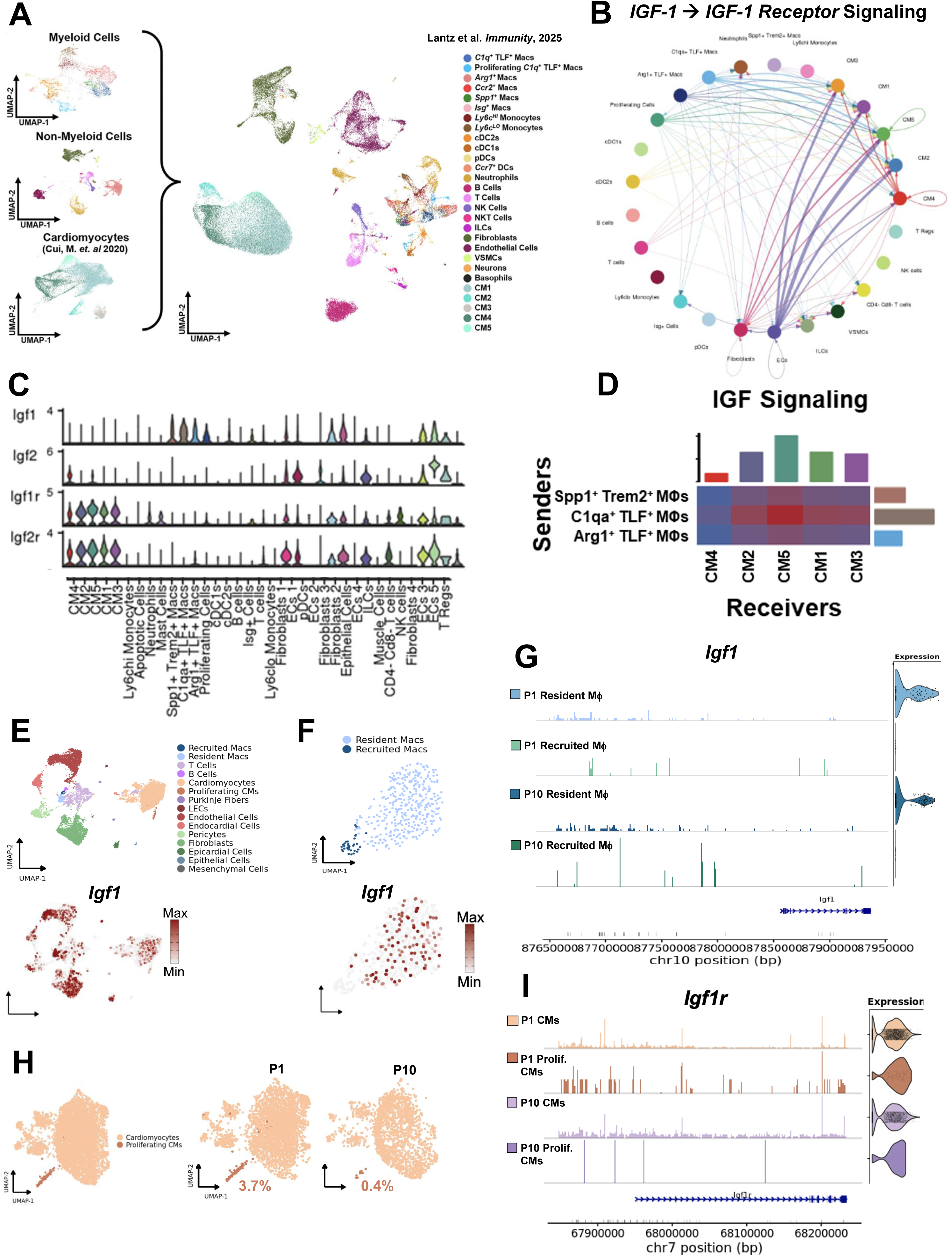
IGF-1 signaling occurs between neonatal *C1q*^+^TLF^+^ *Hif1α*^+^ cardiac macrophages and cardiomyocytes after MI. **A**. Single-nuclei RNA-sequencing from Cui et al. was integrated into our single-cell RNA-sequencing dataset for cell-to-cell communication analyses.^17,19^ B. Circle plots depicting IGF-1 ◊ IGF-1 receptor axes during regeneration. The width of edges represents the strength of the probability of communication. C. Violin plots depicting *Igf1*, *Igf2, Igf1r*, and *Igf2r* expression in neonatal cardiac cell populations 7-days post MI. D. Identification of dominant macrophage IGF-1 senders and cardiomyocyte IGF-1 receivers during neonatal regeneration. E. Expression of *Igf1* across all cardiac cell populations. F. Expression of *Igf1* between resident and recruited cardiac macrophages. G. Chromatic architecture and RNA expression of *Igf1* in resident and recruited macrophages from P1 and P10 hearts. H. Representative proportions of cardiomyocyte and proliferating cardiomyocytes from P1 and P10 hearts. I. Chromatic architecture and RNA expression of *Igf1r* in cardiomyocytes (CMs) and proliferating cardiomyocytes from P1 and P10 hearts Each data point represents a biological replicate from an individual mouse, and data are representative of experiments repeated at least two times. Data with error bars are presented as mean ± SEM. NS= not significant, *, *p* <0.05, ** *p* <0.01, *** *p* <0.001, **** *p* <0.0001.

We then returned to our multiome snRNA-seq and snATAC-seq dataset to identify which P1 and P10 neonatal cardiac cell populations expressed *Igf1* after ischemic cardiac injury (**Figure 5E**). We uncovered that *Igf1* was specifically enriched in resident cardiac macrophages, with loss of *Igf1* message enrichment in recruited macrophages (**Figure 5F**). We also measured distinct differences in *Igf1* chromatin accessibility between both P1 versus P10 recruited and resident cardiac macrophages, which correlated with *Igf1* enrichment in resident cardiac macrophages and loss of *Igf1* enrichment in recruited macrophages (**Figure 5G**). Next, we measured the percentage of proliferating-signature cardiomyocytes in P1 versus P10 neonatal hearts after ischemic cardiac injury and observed an increased percentage of proliferating-type cardiomyocytes in P1 compared to P10 neonates (**Figure 5H**). Lastly, we interrogated differences in P1 versus P10 proliferating cardiomyocyte *Igf1r* chromatin architecture and expression. We identified significant *Igf1r* enrichment in P1 compared to P10 proliferating cardiomyocytes, which correlated with differences in *Igf1r* chromatin accessibility (**Figure 5I**), suggesting that epigenetic regulation of *Igf1r* may contribute to loss of cardiomyocyte proliferative potential. Altogether, our data are consistent with IGF-1 signaling occurring between P1 neonatal *C1q*^+^TLF^+^ *Hif11α^+^* resident cardiac macrophages and cardiomyocytes after ischemic cardiac injury.

### Myocardial IGF-1 infusion rescues cardiac function after MI in *mHif1α* deficient neonatal mice

To determine whether the administration of exogenous IGF-1 improves cardiac function in *mHif1^−/−^* neonatal mice, we injected recombinant IGF-1 (100 ng/mL) into the myocardial wall immediately prior to permanent suture ligation of the LAD coronary artery on P1 (**Figure 6A**). Neonatal *mHif1^−/−^* mice that did not receive myocardial IGF-1 injection served as controls. Twenty-one days post-MI, we measured left ventricular function and examined histologic morphology to assess impact on regeneration. Neonatal *mHif1^−/−^*mice that received IGF-1 injection had significantly reduced myocardial scarring 21 days post-MI compared to *mHif1^−/−^* mice that did not receive IGF-1 injection (**Figure 6B**). Furthermore, left ventricular function was restored in *mHif1^−/−^* mice that received IGF-1 rescue with ejection fraction and fractional shortening values similar to *mHif1^+/+^* neonatal mice 21 days post-MI (**Figure 6C**). Together, our data argue for a molecular mechanism by which P1 neonatal resident cardiac macrophages facilitate cardiac regeneration after ischemic injury through HIF1α transcriptional activation of *Igf1* to modulate neighboring cardiomyocytes for proliferation through IGF-1 signaling (**Figure S5**).

**Figure 6:**
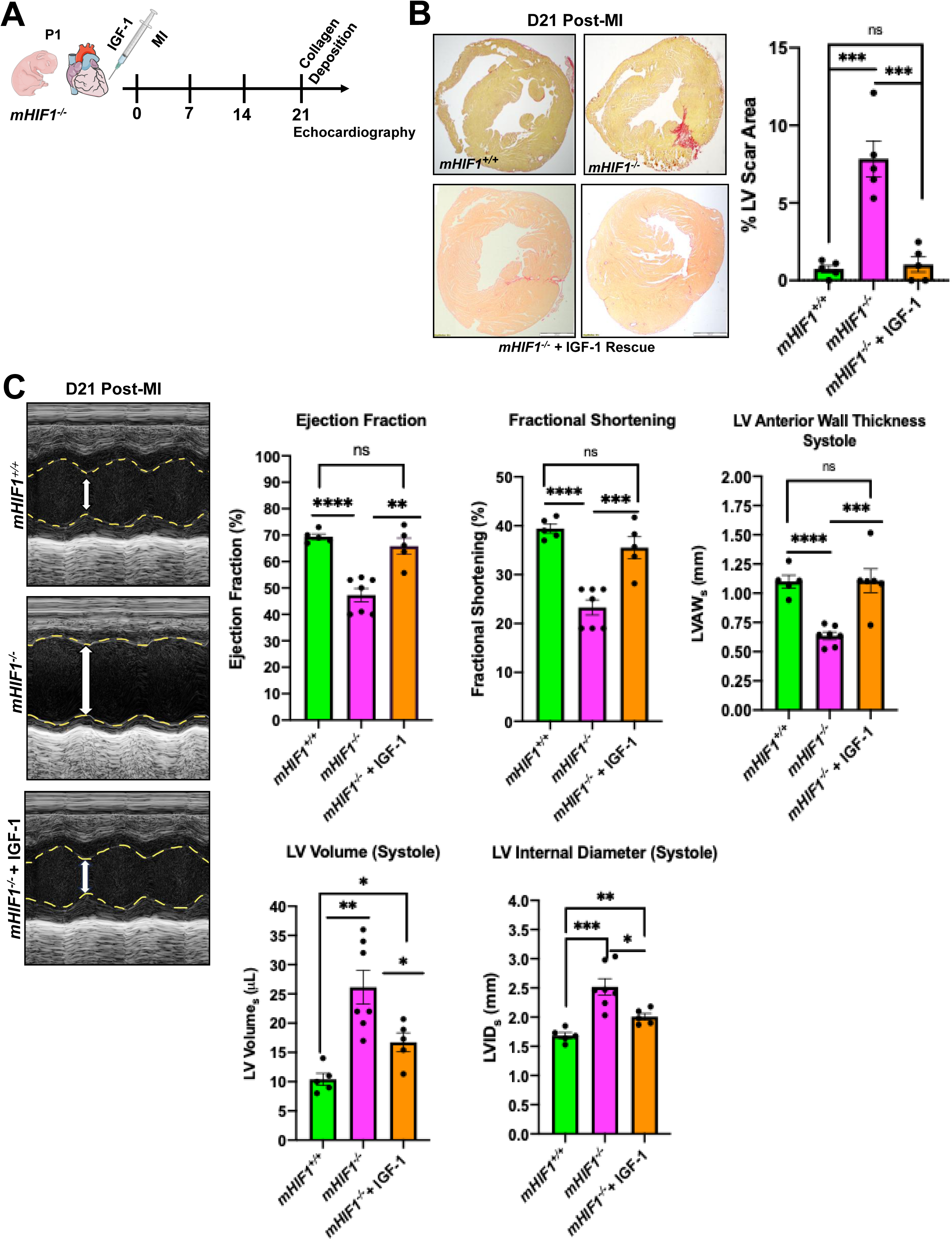
Myocardial IGF-1 injection rescues cardiac function after MI in neonatal mice with genetic depletion of myeloid *Hif1α*. A. Neonatal mice with genetic depletion of *Hif1α* from macrophages (*mHif1^−/−^*) were subjected to permanent LAD ligation to induce MI on P1. Experimental mice were injected with 100 ng/mL of recombinant IGF-1 into the myocardial wall immediately prior to suture ligation of the LAD coronary artery. *mHif1^−/−^* that did not receive IGF-1 injection served as controls. Cardiac function assessed by echocardiography and histology to assess for fibrosis were performed 21-days post MI. B. Assessment of fibrotic scaring using picrosirius red staining of collagen deposition at the site of myocardial injury 21-days post MI in *mHif1^−/−^* + IGF-1 rescue vs *mHif1^−/−^* neonatal mice (*n* = 5). C. Quantification of ejection fraction (EF), fractional shortening (FS), left ventricle anterior wall thickness during systole (LVAW_s_), left ventricle volume during systole, and left ventricle internal diameter during systole (LVID_s_) 21-days post MI in in *mHif1^−/−^* + IGF-1 rescue vs *mHif1^−/−^* neonatal mice (*n* = 5-8). Each data point represents a biological replicate from an individual mouse, and data are representative of experiments repeated at least three times. Data with error bars are presented as mean ± SEM. NS= not significant, *, *p* <0.05, ** *p* <0.01, *** *p* <0.001, **** *p* <0.0001.

## Discussion

Collectively, our findings uncover evidence that macrophage HIF1α facilitates cardiac regeneration in P1 neonatal mice through IGF-1 signaling. With genetic depletion of *Hif1α*, the neonatal regenerative response to ischemic cardiac injury is lost and resembles that of the adult heart with the development of myocardial fibrosis and impaired ventricular function. Specifically, in P1 neonatal mice, HIF1α is stabilized in *C1q*^+^TLF^+^ resident cardiac macrophages after cardiac ischemic injury. Hypoxia stimulates neonatal macrophages to secrete IGF-1, a cardiomyocyte mitogen, through the transcriptional activation of *Igf1* by HIF1α. IGF-1 signaling appears to occur between P1 *C1q*^+^TLF^+^ neonatal cardiac macrophages and cardiomyocytes after MI to support cardiomyocyte proliferation, improve cardiac function, and inhibit myocardial scarring after MI. Furthermore, we uncovered that resident cardiac macrophage function changes in response to cardiac ischemia shortly after birth. Together, the data support a *Hif1α*−dependent mechanism by which tissue hypoxia in the setting of cardiac ischemic injury programs P1 neonatal macrophages to facilitate regeneration.

Macrophage immune responses to ischemic myocardial injury are divergent in neonatal versus adult mice.^20,21^ Ischemic cardiac injury in neonatal mice induces a regenerative response that is dependent on embryonic-derived tissue resident cardiac macrophages.^6,20,21,22^ These macrophages, in contrast to some monocyte-derived macrophages, have been shown to generate minimal inflammation and promote cardiomyocyte proliferation and angiogenesis after cardiac injury.^3,6^ Overall, the underlying mechanisms by which tissue resident cardiac macrophages facilitate cardiac regeneration remain incomplete. Previous studies have identified that neonatal macrophages promote revascularization of the injured myocardium as well as remodeling of the extracellular matrix.^4,23,24^ Regenerative neonatal macrophages have been shown to upregulate C-C motif chemokine ligand 24 (CCL24) and cardiotrophin-like cytokine 1 (CLCF1) which can induce neonatal cardiomyocyte proliferation *in vitro*.^25,26^ Recently, we discovered that MerTK directed efferocytosis remodels arachidonic acid metabolism in neonatal *C1q*^+^TLF^+^ resident cardiac macrophages to facilitate cardiac regeneration through the generation of thromboxane A_2_ (TXA_2_).^17^ In the future, it will be informative to determine whether HIF1α and IGF-1 work in synergy with TXA_2_ to initiate and enhance cardiac regeneration.

Hypoxia is a molecular stimulus for fetal development and regenerative processes.^27^ In neonatal mice, hypoxia has been linked to the proliferation of embryonic-derived tissue resident cardiac macrophages.^28^ These macrophages facilitate not only cardiac regeneration but also fetal cardiogenesis, processes that both occur in hypoxic environments.^21,29,30^ In neonatal mice, hypoxia has been linked to increased cardiomyocyte proliferation with cardiomyocyte proliferation dependent on resident cardiac macrophages.^28^ In utero, both murine and human hearts are subjected to low O_2_ concentrations, ranging from 1-8% depending on gestational age.^31,32,33,34^ In response to hypoxia, fetal macrophages favor expression of *Hif1α*, as evidenced by human neonatal cord blood macrophages expressing higher levels of *Hif1α* than adult macrophages.^3,35^ In utero, murine macrophage HIF1α regulates cardiac development and fetal survival.^36,37^ HIF1α also coordinates a transcriptional program that prioritizes glycolysis.^38,39,40,41^ With increased oxygen availability at birth, myocardial metabolism shifts from glycolysis to oxidative phosphorylation with subsequent loss of cardiomyocyte replication and cell cycle arrest due to oxidate DNA damage.^3,42^ The shift in these metabolic states coincides with the loss of macrophage regenerative function and the short regenerative window of the neonatal heart.^3,17,35^ Why this occurs remains a critical knowledge gap. Cell culture studies using human cancer cells have identified that hypoxia induces specific changes in chromatin accessibility which correlate with changes in gene expression.^43^ Furthermore, most hypoxia-induced changes to chromatin accessibility were HIF-dependent and reversible upon reoxygenation.^43^ Whether perinatal changes in oxygenation or metabolism induce changes in chromatin accessibility, *Hif1α* expression, or HIF1α regenerative target genes that result in loss of macrophage regenerative function is unknown. Future interrogation of the epigenetic regulation of HIF1α and regenerative targets during perinatal transition warrants further research.

We have newly identified that hypoxia stimulates HIF1α transcriptional activation of *Igf1* in neonatal resident cardiac macrophages with subsequent release of IGF-1, stimulating cardiomyocyte proliferation. IGF-1 has previously been shown to play an important role in fetal development and early postnatal life. Specifically, IGF-1 supports overall fetal survival, fetal and neonatal growth, central nervous system development, and lung development.^44^ In non-mammalian species, such as zebrafish, inhibition of IGF-1 receptor signaling has been shown to block cardiomyocyte proliferation during both heart development and cardiac regeneration.^45^ In the setting of adult murine hypertensive cardiac stress, resident cardiac macrophages secrete IGF-1 to facilitate cardiomyocyte growth and adaptive remodeling to preserve cardiac function.^46^ In adult murine MI, IGF-1 has been shown to reduce scarring and improve cardiac function after MI by polarizing macrophages and neutrophils to anti-inflammatory phenotypes.^47^ Our study newly adds understanding to the role of IGF-1 in mammalian neonatal cardiac regeneration.

Our study is not without limitations. For example, we cannot rule out the contribution of *Hif1α* from other myeloid cells or extra cardiac sources on cardiac regeneration. It also remains to be determined the effect of silencing IGF-1 specifically from cardiac macrophages *in vivo*. Furthermore, we employed transcriptomic data to statistically infer IGF-1 signaling between neonatal *C1q*^+^TLF^+^ cardiac macrophages and cardiomyocytes *in vivo* after MI. We also utilized peripheral neonatal macrophages rather than resident cardiac macrophages to interrogate *Hif1α* mediated regenerative mechanisms *in vitro*. Although this remains an important limitation of our study, it is notable that HIF1α regenerative potential is also conserved in neonatal peripheral macrophages.

In conclusion, we have newly demonstrated that macrophage *Hif1α* is essential for neonatal cardiac regeneration after MI through the transcriptional activation of *Igf1* and subsequent IGF-1 signaling between resident cardiac macrophages and cardiomyocytes. In future studies, it will be interesting to interrogate the epigenetic regulation of *Hif1α* and regenerative targets to uncover how and when *Hif1α*-mediated macrophage regenerative function is lost.

## Novelty and Significance

### What is Known?

- Neonatal mice, in contrast to adult mice and humans, can readily regenerate injured myocardium after ischemic injury.
- Regenerative capacity is dependent on macrophages and lost shortly after birth.

### What new information does this article contribute?

- Distinct neonatal cardiac macrophage populations respond uniquely to ischemic injury, with resident cardiac macrophage function changing shortly after birth.
- Myeloid *Hif1α* is necessary for neonatal cardiac regeneration after ischemic cardiac injury.
- *Hif1α* facilitates cardiac regeneration through the transcriptional activation of *Igf1* with subsequent IGF-1 signaling between neonatal resident cardiac macrophages and cardiomyocytes post-MI.

### Summary of Novelty and Significance

Neonatal mice, in contrast to adult mice, possess the innate capacity to regenerate injured myocardium after myocardial infarction. Neonatal regenerative capacity is dependent on resident cardiac macrophages. We have newly identified that myeloid *Hif1α* is essential for neonatal cardiac regeneration after MI through the transcriptional activation of *Igf1* in resident cardiac macrophages and subsequent IGF-1 signaling with neonatal cardiomyocytes.

## Acknowledgments

These studies employed sequencing services provided by the NUSeq Core at Northwestern University. This research was also supported in part through the computational resources and staff contributions provided by the Quest high performance computing facility at Northwestern University, which is jointly supported by the Office of the Provost, the Office for Research, and Northwestern University Information Technology.

## Author Contributions

Conceptualization, A.B., C.L., and E.B.T.; Methodology, A.B., C.L., and Z-D.G.; Mice, M.D., and E.B.T; Investigation: A.B., C.L., A.A., and K.G.; Formal Analysis, A.B. and C.L.; Software, A.B. and C.L.; Resources, A.B. and E.B.T.; Writing Original Draft, A.B. and E.B.T.; Writing Review and Editing, all authors; Funding Acquisition, A.B. and E.B.T.; Supervision, E.B.T.

## Non-standard Abbreviations and Acronyms

HIF1α: Hypoxia-inducible transcription factor 1 alpha
IGF-1: Insulin-like growth factor 1
MI: Myocardial infarction
BMDMs: Bone marrow-derived macrophages
IGF1R: Insulin-like growth factor 1 receptor

## Supplemental Figures

Please see supplemental information.

